# Abl kinases regulate FGF signaling independent of Crk phosphorylation to prevent Peters anomaly

**DOI:** 10.1101/2024.10.24.619064

**Authors:** Hao Wu, Yingyu Mao, Qian Wang, Honglian Yu, Michael Bouaziz, Neoklis Makrides, Anthony J. Koleske, Glenn L. Radice, Xin Zhang

## Abstract

Peters anomaly, the most common cause of congenital corneal opacity, stems from corneal-lenticular adhesion. Despite numerous identified mutations, a cohesive molecular framework of the disease’s etiology remains elusive. Here, we identified Abl kinases as pivotal regulators of FGF signaling, as genetic ablation of Abl kinases restores lens induction even in the absence of FGF signaling. Intriguingly, both *Abl* kinase deficiency and increased FGF-Ras activity result in Peters anomaly independent of ERK signaling, which can be rescued by allelic deletion of Abl substrate, Crk. However, contrary to the prevailing belief that Abl kinases regulate Crk proteins by direct phosphorylation, mutations at Abl kinase phosphorylation sites on Crk and CrkL did not yield any observable effects. Instead, our findings reveal that Abl kinases phosphorylate Ptpn12, which in turn inhibits p130Cas phosphorylation and Crk recruitment, crucial for Rho GTPases activation and cytoskeletal dynamics. Consequently, Abl kinase deficiency reduces actomyosin contractility within the lens vesicle and genetically interacts with RhoA inhibition. Conversely, *Rac1* deletion mitigates Peters anomaly in models with aberrant FGF, Abl kinase and RhoA signaling. Our results demonstrate that Abl kinases regulate FGF signaling to balance RhoA and Rac1 activity via the Ptpn12-p130Cas pathway, suggesting that targeting tension-mediated lens vesicle separation could be a therapeutic strategy for Peters anomaly.

## Introduction

Peters anomaly is a congenital disorder that primarily affects the anterior segment of the eye, leading to various vision impairments, including central corneal opacity, iridocorneal adhesions, and congenital glaucoma (Bhandari et al., 2011). It has been linked to several genes, mainly transcription factors -including *PAX6*, *PITX2*, *MAF*, and *FOXC1*, although the specific molecular pathways involved remain elusive. A notable feature of Peters anomaly is the development of corneal-lenticular adhesions, which stem from the failure of the lens vesicle to separate from the surface ectoderm during embryonic development, leaving a residual lens stalk behind (Cvekl and Zhang, 2017). Intriguingly, the lens vesicle arises from the surface ectoderm that expresses both E-cadherin and P-cadherin, but it turns on the expression of N-cadherin at the expense of P-cadherin (Xu et al., 2002). This cadherin switch is thought to facilitate cell sorting at this developmental stage, contributing to the segregation of the lens vesicle from the surface ectoderm, but this model remains to be tested.

FGF signaling has also been associated with Peters anomaly. In humans, sporadic cases of Peters anomaly have been noted in patients with gain-of-function mutations in *FGFR2*, including some with Crouzon or Pfeiffer syndromes (McCann et al., 2005; Okajima et al., 1999). Supporting this, mouse studies have demonstrated that either deleting *Spry*, a negative regulator of FGF signaling, or overexpressing FGF or Ras, can lead to the formation of a persistent lens stalk (Burgess et al., 2010; Kuracha et al., 2011; Thein et al., 2016). During lens development, FGF signaling exhibits a dynamic pattern of activity, both spatially and temporally, as evidenced by the expression of phospho-ERK (pERK) (Makrides et al., 2022). Initially, pERK activity is low in the surface ectoderm but increases as it thickens to form the lens placode. As the placode invaginates to become the lens vesicle, pERK remains elevated within the interior of the vesicle but decreases significantly in the lip regions connected to the surface ectoderm. How this precise pattern of FGF signaling regulates the timely separation of the lens from the surface ectoderm remains to be understood.

The tyrosine kinase ABL was first discovered as an oncoprotein in human leukemia, where it promotes many aspects of cellular transformation, including increased cell proliferation, survival, and invasion (Bradley and Koleske, 2009; Greuber et al., 2013). A landmark achievement in targeted therapy for cancer has been the development of Imatinib (Gleevec), a small molecule that inhibits the BCR-ABL fusion protein. Genetic studies in mice have also revealed the critical importance of *Abl* kinase genes in embryonic development. While *Abl* knockout mice die around birth and *Arg* (a close homolog) null mutants are viable, combined deletion of both *Abl* and *Arg* results in early embryonic lethality at E8 (Koleske et al., 1998; Schwartzberg et al., 1991; Tybulewicz et al., 1991). One of the key substrates of Abl kinases is Crk, which was also originally discovered as the cellular counterpart of the viral oncoprotein v-Crk from avian sarcoma retrovirus (Birge et al., 2009). Unlike Abl kinases, however, Crk is non-catalytic, consisting of an SH2 domain and two SH3 domains, thus the transforming activities of v-Crk and Crk are attributed to their unfettered function as adaptor proteins. Biochemical and structural studies have shown that Abl kinases phosphorylate a conserved tyrosine residue between two SH3 domains of Crk and its homolog CrkL, leading to the intramolecular binding by SH2 and sequestration of both the SH2 and N-SH3 domains (Cho and Klemke, 2000; Huang et al., 2008; Jankowski et al., 2012; Saleh et al., 2016). This elegant regulatory mechanism presents an eminent example of how protein tyrosine kinases can control signal transmission by modulating the accessibility of adaptor proteins.

In this study, we showed that mice lacking *Abl* and *Arg* genes develop Peters anomaly. Although *Abl/Arg* deficiency rescued the lens development failure in FGF signaling mutants by elevating ERK activity, it led to the persistent lens stalk due to heightened Crk signaling. Unexpectedly, mutations in Abl kinase phosphorylation sites on Crk and CrkL proteins did not produce noticeable effects. Instead, our analyses showed that Abl kinases reduce Crk signaling by stabilizing Ptpn12, which in turn dephosphorylates the Crk-binding protein p130Cas. At the cellular level, although *Abl/Arg* deficiency destabilizes adherens junction in vitro, ablation of E-cadherin does not aggravate the *Abl/Arg* phenotype. Contrary to the cadherin switch model, eliminating both P-and N-cadherin also failed to induce a lens stalk phenotype. Instead, we observed that Abl kinases promote actinomyosin contractility in the lens vesicle by shifting the Rac1-RhoA antagonism. Supporting this, *Rac1* deletion alleviates the lens stalk abnormality caused by the reduced Abl kinases and RhoA activities or increased FGF-Ras signaling. These results demonstrated that Abl kinases oppose FGF signaling through the Ptpn12-p130Cas axis, promoting the tension-mediated lens vesicle separation from the surface ectoderm.

## Results

### The lens-specific ablation of Abl kinases causes Peters anomaly

We generated lens-specific *Abl* and *Arg* knockout mice by crossing floxed alleles with *Le-Cre*, which is specifically active in the lens starting at E9.5 (Ashery-Padan et al., 2000). The Cre-associated GFP reporter revealed a unique ring structure in the center of mutant eyes (Fig. 1A, yellow arrow), identified through histological analysis as corneal-lenticular adhesions (Fig. 1A, black arrow). Unlike control animals with clear iris-cornea separation, *Abl/Arg* mutants consistently showed iridocorneal adhesions (Fig. 1A, arrowheads), resembling human Peters anomaly. Further embryonic development analysis confirmed *Le-Cre* activity (indicated by GFP expression) was confined to the lens vesicle at E10.5, but mutant lens did not display any changes in cell proliferation as shown by phospho-histone3 (pHH3) staining (Fig. 1B and C). Similarly, TUNEL assay and cleaved caspase 3 staining revealed normal cell apoptosis patterns at the rim of the invaginating lens vesicles (Fig. 1D and E, arrowheads). By E12.5 when the control lens was completely detached from the surface ectoderm, *Abl/Arg* mutant displayed persistent lens stalks consisting of Pcad-expressing surface ectoderm extending into Ncad-expressing lens epithelium (Fig. 1F, arrow). This prevented the proper closure of the lens vesicle, evident from gaps in Foxe3 (lens epithelial marker) and Col IV (lens capsule marker) expression (Fig. 1F, arrowheads). At E14.5, *Abl* and *Arg* loss did not impact cell proliferation or cell cycle exit (shown by Ki67 and p57 staining) or prevent expression of transcription factors p63 in the surface ectoderm and Pax6, Foxe3, and Maf in the lens (Fig. 1G and H). However, the mutant lens remained open, occasionally allowing Jag1-expressing lens fibers to protrude from the eye (Fig 1J, arrow). These findings demonstrate that Abl family kinase deletion causes Peters anomaly in mice without compromising lens cell differentiation, survival, or proliferation.

**Figure 1.**
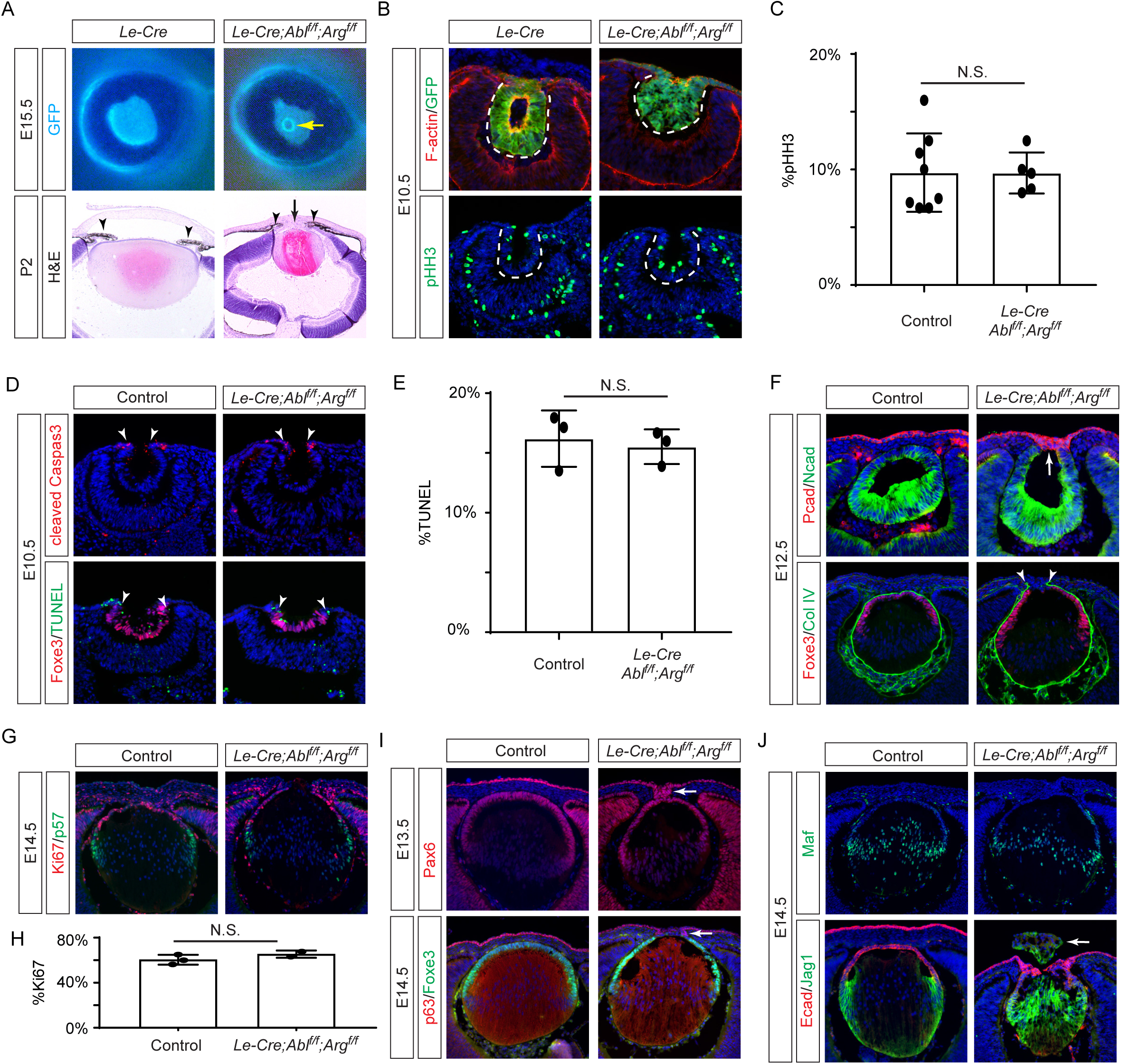
Genetic Ablation of *Abl/Arg* Results in Peters Anomaly in Mice. **(A)** *Abl/Arg* mutants exhibited persistent lens stalks visible as a ring structure illuminated by the GFP reporter from the *Le-Cre* driver (yellow arrow). P2 mutant animals showed the aberrant iridocorneal (arrowheads) and cornea-lenticular adhesion (black arrows) characteristic of Peters anomaly. **(B)** The *Le-Cre* driver is specific expressed in the lens vesicle as shown by the companying GFP reporter, but deletion of *Abl/Arg* did not affect cell proliferation shown by pHH3 staining. **(C)** Quantification of the percentage of pHH3-positive cell in the lens vesicle. Student’s t-test, n=8 for control, n=5 for *Abl/Arg* mutants, N.S., Not Significant. **(D)** *Abl/Arg* deletion did not affect cell apoptosis as shown by cleaved caspase3 and TUNEL staining (arrowheads). **(E)** Quantification of the percentage of TUNEL-positive cells in the lens vesicle. Student’s t-test, n=3, N.S., Not Significant. **(F)** In *Abl/Arg* mutants, the Ncad-positive lens vesicle failed to close, as evidenced by ColIV staining (arrowheads), and remained attached to the Pcad-positive surface ectoderm (arrow). **(G)** Cell proliferation and cell cycle exit were unaffected in E14.5 *Abl/Arg* mutants as shown by Ki67 and p57 staining. **(H)** Quantification of the percentage of Ki67-positive cells in the lens. Student’s t-test, n=3 for control, n=2 for *Abl/Arg* mutants, N.S., Not Significant. **(I)** Despite the persistent lens stalks (arrows), the distinct expression patterns of p63 in the surface ectoderm and Foxe3 in the lens vesicle were maintained in *Abl/Arg* mutants, along with co-expression of Pax6. **(J)** While lens fiber differentiation, indicated by expression of Maf and Jag1, occurred properly, severe *Abl/Arg* mutants exhibited lens herniation (arrow).

### Abl kinases inhibit FGF-ERK signaling in lens induction

Previous studies have implicated hyperactive FGF signaling in Peters anomaly (Burgess *et al*., 2010; Kuracha *et al*., 2011; McCann *et al*., 2005; Okajima *et al*., 1999; Thein *et al*., 2016). Our experiments further substantiate this by demonstrating that overexpressing *Fgf3* in the lens (*Tg-Fgf3*) or activating an inducible allele of oncogenic Kras (*Kras ^G12D^*) by *Le-Cre* both resulted in a persistent lens stalk (Fig. 2A, arrowheads) (Robinson et al., 1998). Additionally, the *Tg-Fgf3* mutant displayed extrusion of the lens materials outside the eye, akin to what is observed in *Arg/Abl* mutants (Fig. 2A, arrows). Given the phenotypic similarities between *Abl/Arg* knockouts and FGF-Ras gain-of-function mutants, we explored a possible connection between Abl kinases and FGF signaling. We infected *Arg^f/f^;Abl^f/f^* MEF cells with a Cre-expressing virus (Ad-Cre), effectively ablating Abl/Arg and reducing the phosphorylation of its substrate, Crk. Interestingly, this resulted in an increase in pERK levels compared to MEF cells infected with a GFP-expressing virus (Ad-GFP) (Fig. 2B). This was further confirmed by acute inhibition of Abl kinase activity by pharmacological inhibitors. Unlike the dual Src/Abl inhibitor Dasatinib, which reduced both pCrk and pERK levels, the Abl-specific inhibitor Imatinib decreased pCrk but increased pERK. Lastly, we showed that the pERK level was elevated in *Arg/Abl* mutant lens (Fig. 2C), demonstrating that Abl kinases negatively regulate ERK signaling during lens development.

**Figure 2.**
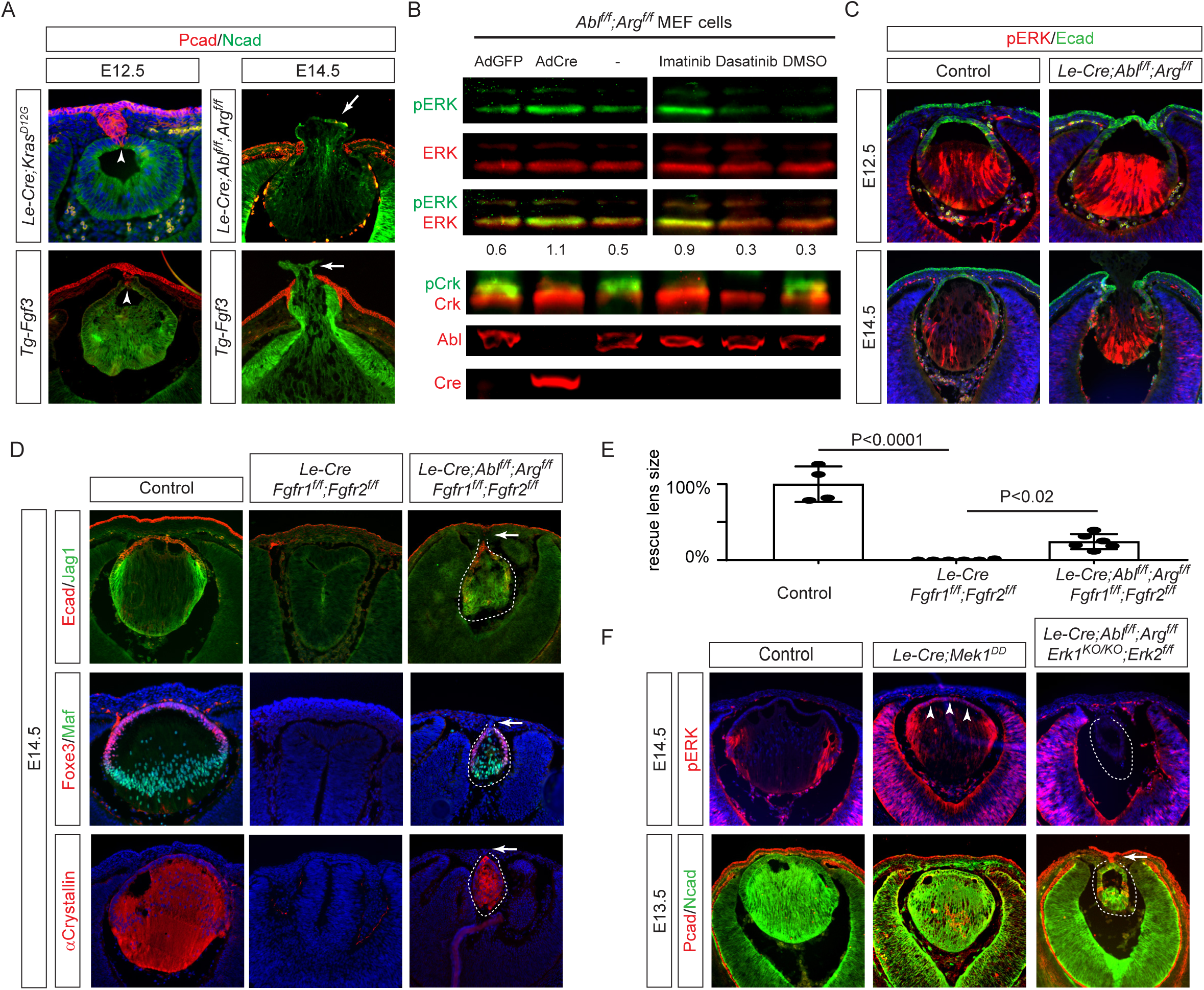
*Abl/Arg* deficiency Mimics Increased FGF Signaling in Peters Anomaly and Lens Induction. **(A)** Expression of *Fgf3* or constitutively active *Kras^G12D^* resulted in persistent lens stalk (arrowheads) and lens herniation (arrows) similar to that observed in *Abl/Arg* mutant. **(B)** Treatment with the dual Src/*Abl* inhibitor Dasatinib eliminated both pERK and pCrk signaling, whereas the *Abl*-specific inhibitor Imatinib inhibited pCrk but enhanced pERK levels. Genetic ablation of *Abl/Arg* in MEF cells using an Ad-Cre (adenovirus expressing Cre recombinase) resulted in a loss of pCrk and an increase in pERK, unlike control cells infected with an Ad-GFP (adenovirus expressing GFP), where these levels remained unchanged. **(C)** *Abl/Arg* mutant lens exhibited increased pERK staining. **(D)** Lens development was abolished in *Fgfr1/2* mutant, but was partially restored after *Abl/Arg* deletion, although corneal-lenticular adhesions persisted (arrows). **(E)** Quantification of the lens size. One way ANOVA, n=4 for control, n=6 for *Fgfr1/2* and *Fgfr1/2/Abl/Arg* mutants, *P<0.001, **P<0.02. **(F)** Expression of constitutively active *Mek1^DD^* expanded pERK expression to the anterior lens epithelium but did not result in lens stalk formation. Conversely, deletion of *Erk1/2* did not rescue the incomplete lens vesicle separation defect observed in *Abl/Arg* mutants.

We further explored the genetic interaction between *AblArg* and FGF signaling. As we and others have previously shown, genetic ablation of *Fgfr1* and *Fgfr2* abolished lens development (Collins et al., 2018; Garcia et al., 2011). Remarkably, further deletion of *Abl/Arg* in *Fgfr1/2* mutants resulted in the formation of a well-structured lens, characterized by Foxe3-positive lens epithelium and lens fibers expressing Maf, Jag1, and αCrystallin (Fig. 2D, dotted lines). Despite this, the hypoplastic lens remained attached to the ectoderm. To further explore the role of MAPK signaling in Peters anomaly, we crossed *Le-Cre* with the Cre-inducible *Mek1 ^DD^* allele to activate Mek1 constitutively in the lens (Srinivasan et al., 2009). This led to an expansion of pERK activity from the lens equator to the anterior epithelium, yet did not lead to the formation of a lens stalk (Fig. 2F, arrowheads). Additionally, combining *Erk1/2* knockout with *Abl/Arg* deletion significantly reduced lens size, but failed to prevent persistent lens-cornea attachment (Fig. 2F, arrow). These findings indicate that while Abl kinases negatively regulate FGF signaling by inhibiting MAPK signaling during lens induction, the deficiency in Abl kinase signaling contributes to Peters anomaly independently of its effects on ERK signaling.

### Crk proteins mediate FGF and Abl kinase signaling in lens vesicle separation independent of tyrosine phosphorylation

Abl kinases phosphorylate Crk and CrkL at Y221 and Y207 in the linker regions, respectively, which competes with the binding partners of the SH2 domains of Crk and CrkL to terminate signaling (Cho and Klemke, 2000; Huang *et al*., 2008; Jankowski *et al*., 2012; Saleh *et al*., 2016). Indeed, we found that pCrk was abolished in *Abl/Arg* mutant lens vesicles, confirming Abl and Arg as the primary kinases responsible for Crk protein phosphorylation (Fig. 3A, dotted lines). We investigated the genetic interactions between *Crk/CrkL* and *Abl/Arg* in lens development. Notably, genetic ablation of both *Crk* and *CrkL* reversed the increase in pERK observed in *Abl/Arg* mutants and rescued the lens vesicle separation phenotype, although it also reduced lens size (Fig. 3B). We generated an allelic series of *Crk* and *CrkL* deletion and found deleting at least two of the four alleles effectively prevented lens stalk formation in *Abl/Arg* mutants without significantly impacting lens size. (Fig. 3C, D and F). Removing only one *CrkL* allele, however, was insufficient to ameliorate the lens stalk phenotype (Fig. 3D, arrow), indicating a dose-dependent effect of Abl-Crk signaling on lens vesicle separation. Additionally, deleting *Crk/CrkL* in the *Tg-Fgf3* mutant abrogated the lens stalk phenotype (Fig. 3E, arrow), providing strong evidence that Crk mediates both FGF and Abl kinase signaling pathways during lens vesicle separation.

**Figure 3.**
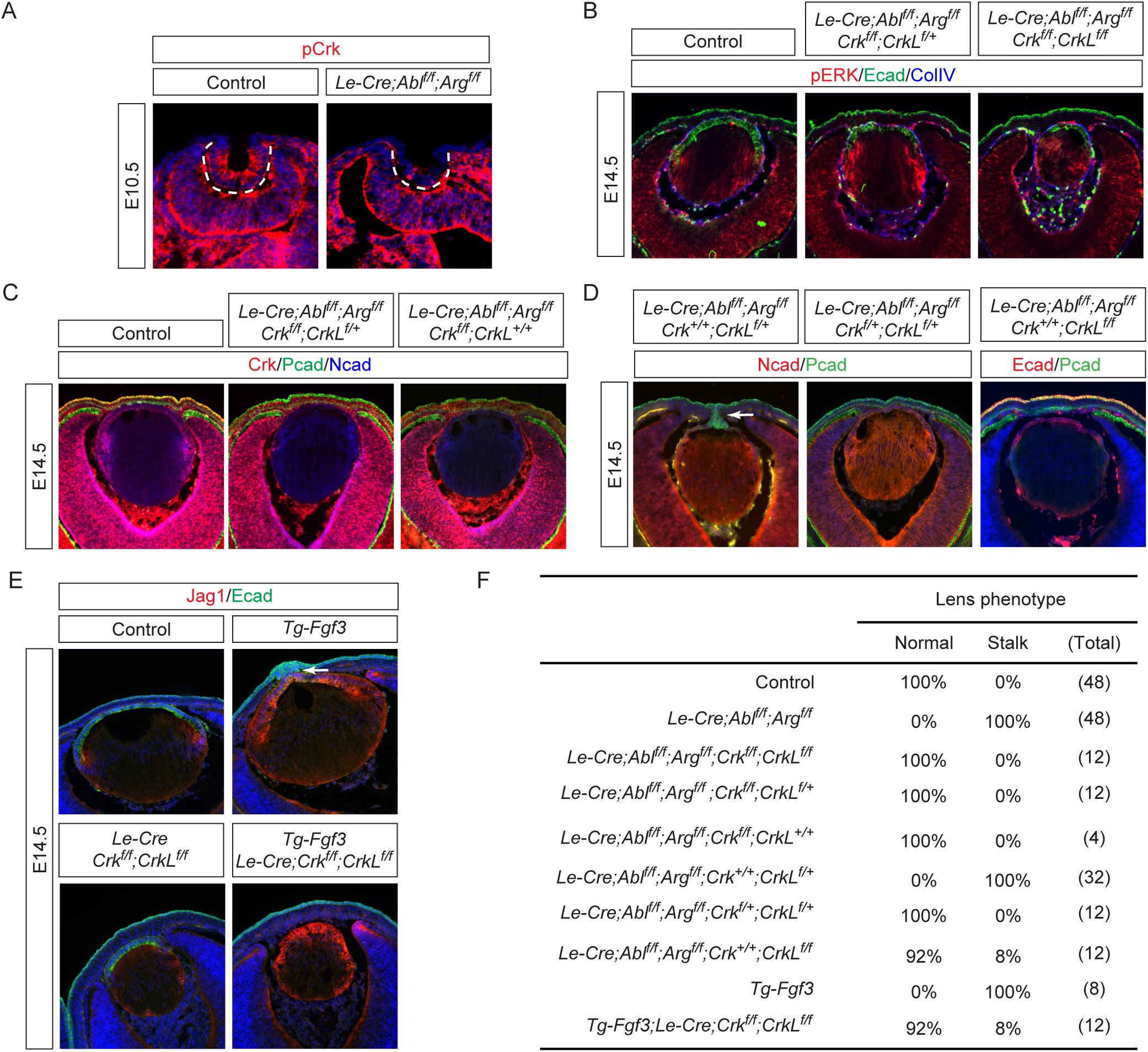
Attenuation of Crk Signaling Restores Lens Vesicle Separation in Both *Abl/Arg* and FGF Mutants. **(A)** pCrk was lost in *Abl/Arg* mutant lens vesicle (dotted lines). **(B)** Complete deletion of both *Crk* and *CrkL* in *Abl/Arg* mutants reduced pERK expression and eliminated the lens stalk phenotype. **(C)** Deletion of Crk alone, evident from the loss of Crk staining, was enough to facilitate the separation of the lens vesicle from the surface ectoderm in *Abl/Arg* mutants. **(D)** Amelioration of the lens stalk phenotype in *Abl/Arg* mutants was achieved by removing two or three alleles of Crk and CrkL, but not with the removal of only one allele (arrow). **(E)** *Crk* and *CrkL* deletion prevented the lens-cornea attachment in the *Tg-Fgf3* mutant (arrow). **(F)** Quantification of the lens stalk phenotype.

To directly test the hypothesis that Abl kinase phosphorylation of Crk is necessary for lens vesicle separation, we generated a *Crk* Y221F knock-in mutation (*Crk ^YF^*) to abolish the Abl kinase phosphorylation site (Fig. 4A and B). Western blot analysis revealed that this mutation not only abolished pCrk (Fig. 4C, upper band) but also increased the mobility of unphosphorylated Crk (Fig. 4C, lower band). Crk proteins are known to play a role in integrin-mediated focal adhesion complexes for cell-matrix interactions (Birge *et al*., 2009). Unexpectedly, *Crk ^YF/YF^* MEF cells showed normal focal adhesion maturation as indicated by the progressive increase in phosphorylation of Paxillin when plated on fibronectin-coated dishes. Immunoprecipitation studies also showed that *Crk ^YF^* maintained the same affinity to the upstream protein p130Cas (Fig. 4D). Remarkably, homozygous *Crk ^YF/YF^* animals were viable and fertile, with normal lens vesicle closure and no residual attachment to the surface ectoderm (Fig. 4E). Even when crossed onto a *CrkL* deletion background, the resulting *Le-Cre;Crk ^YF/YF^*;*CrkL ^f/f^*mutants showed no differences in lens size from the single *CrkL* mutants (Fig. 4F and G), suggesting that *CrkL* does not compensate for the *Crk ^YF/YF^* mutant. To further investigate these surprising results, we used CRISPR-Cas9 technology to generate a *CrkL* Y207F mutation (*CrkL^YF^*) (Fig. 4H). Like the *Crk ^YF^* mutation, *CrkL^YF^* also displayed a loss of phosphorylation and a mobility shift in western blots (Fig. 4I). In *Crk ^YF/YF^*;*CrkL^YF/YF^*cells, both proteins appeared as a single unphosphorylated band. However, *Crk ^YF/YF^*;*CrkL^YF/YF^* mutant animals are normal and healthy, and their lenses showed no alterations in pERK levels, marker expression, or size (Fig. 4J and K). These findings decisively demonstrate that, contrary to long-held beliefs, Abl kinase’s phosphorylation of Crk and CrkL is dispensable for their activities.

**Figure 4.**
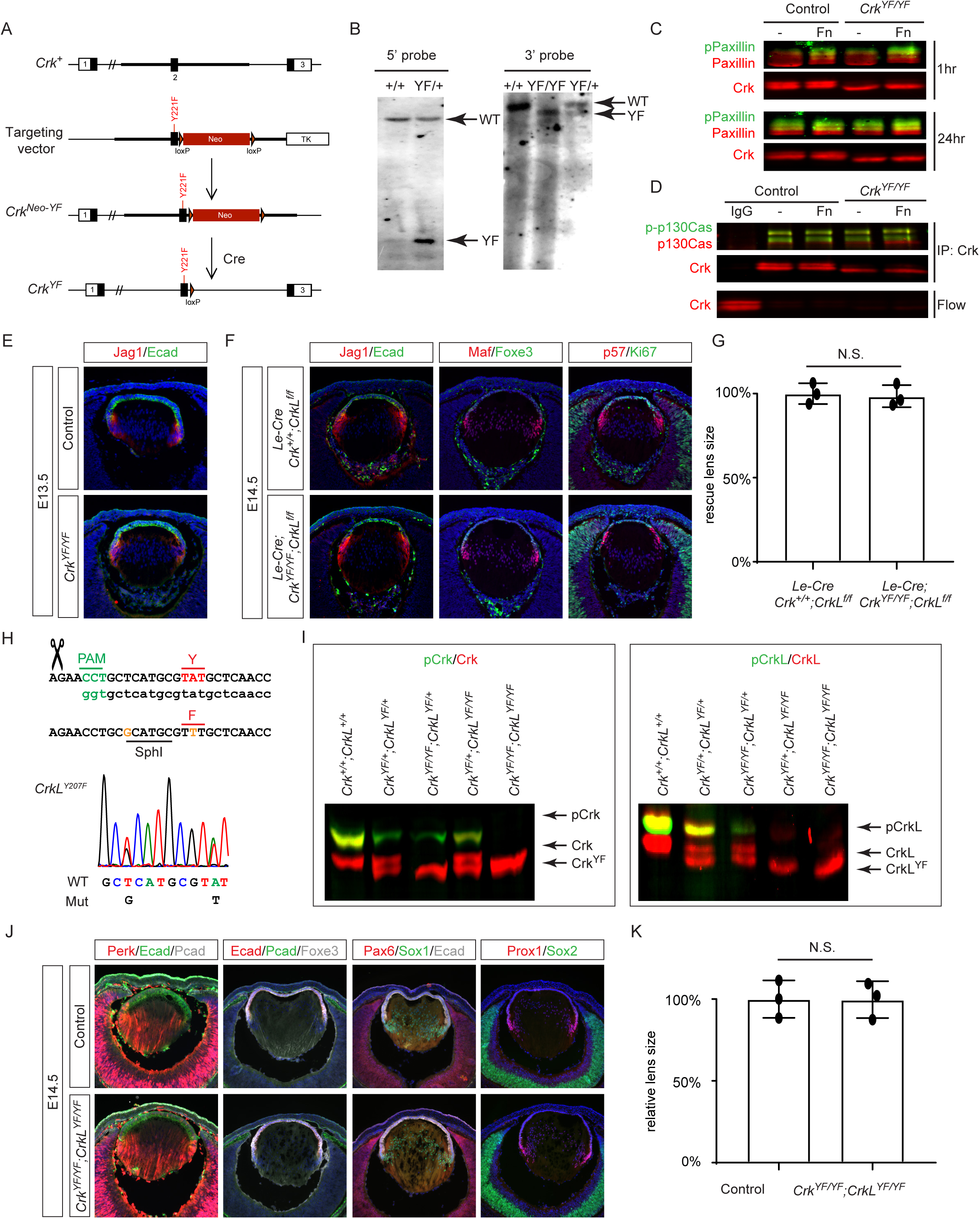
Tyrosine Phosphorylation of Crk and CrkL is Dispensable for Normal Development. **(A)** Schematic of the genetic targeting strategy used to create the Y221F mutation in *Crk* (*Crk^YF^*). **(B)** Validation of the *Crk^YF^* mutation by southern blots. **(C)** Despite the Y221F mutation causing an apparent mobility shift in Crk, *Crk^YF/YF^* MEF cells exhibited a similar increase in pPaxillin levels to control cells when adhered to fibronectin-coated plates over a period of 1 to 24 hours. **(D)** Immunoprecipitation assays demonstrated that Crk^YF^ binds to p130Cas similarly to the wild-type Crk protein. **(E)** *Crk^YF/YF^* lens did not exhibit any noticeable phenotypic abnormalities. **(F)** Even with concurrent deletion of *CrkL*, *Crk^YF^* mutation did not disrupt lens development. **(G)** Quantification of the lens size. One way ANOVA, n=3, N.S., Not Significant. **(H)** Generation of the *CrkL* Y209F mutation (*CrkL^YF^*) by CRISPR-Cas9 technology. **(I)** Western blot analysis confirmed the absence of pCrk and pCrkL in *Crk^YF^*and *CrkL^YF^* mutants. **(J)** *Crk^YF/YF^*;*CrkL^YF/YF^* mutants displayed no lens phenotype. **(K)** Quantification of the lens size. One way ANOVA, n=3, N.S., Not Significant.

### Lens vesicle separation requires the Abl kinase activity but not Vinculin phosphorylation

The lack of phenotype in *Crk ^YF/YF^*;*CrkL^YF/YF^* mutants raises doubt whether the catalytic activity of Abl kinase is necessary for lens vesicle development. To test this idea, we use CRISRP-Cas9 technology to knock in the K270R kinase dead mutation (*Abl^KD^*) in the *Abl* locus, eliminating the catalytic lysine residue in the ATP-binding pocket critical for Abl kinase activity (Hardin et al., 1996) (Fig. 5A). As expected, homozygous *Abl^KD^* mutant mice exhibited perinatal lethality similar to null mutants and pCrk was lost in *Abl ^f/KD^;Arg ^f/f^* MEF cells after Cre-mediated removal of the floxed alleles (Fig. 5B). Notably, *Le-Cre;Abl ^f/KD^;Arg ^f/f^*mutant lens displayed enhanced pERK staining and failed lens vesicle separation (Fig. 5C), demonstrating that Abl kinase activity is indeed essential for normal lens development.

**Figure 5.**
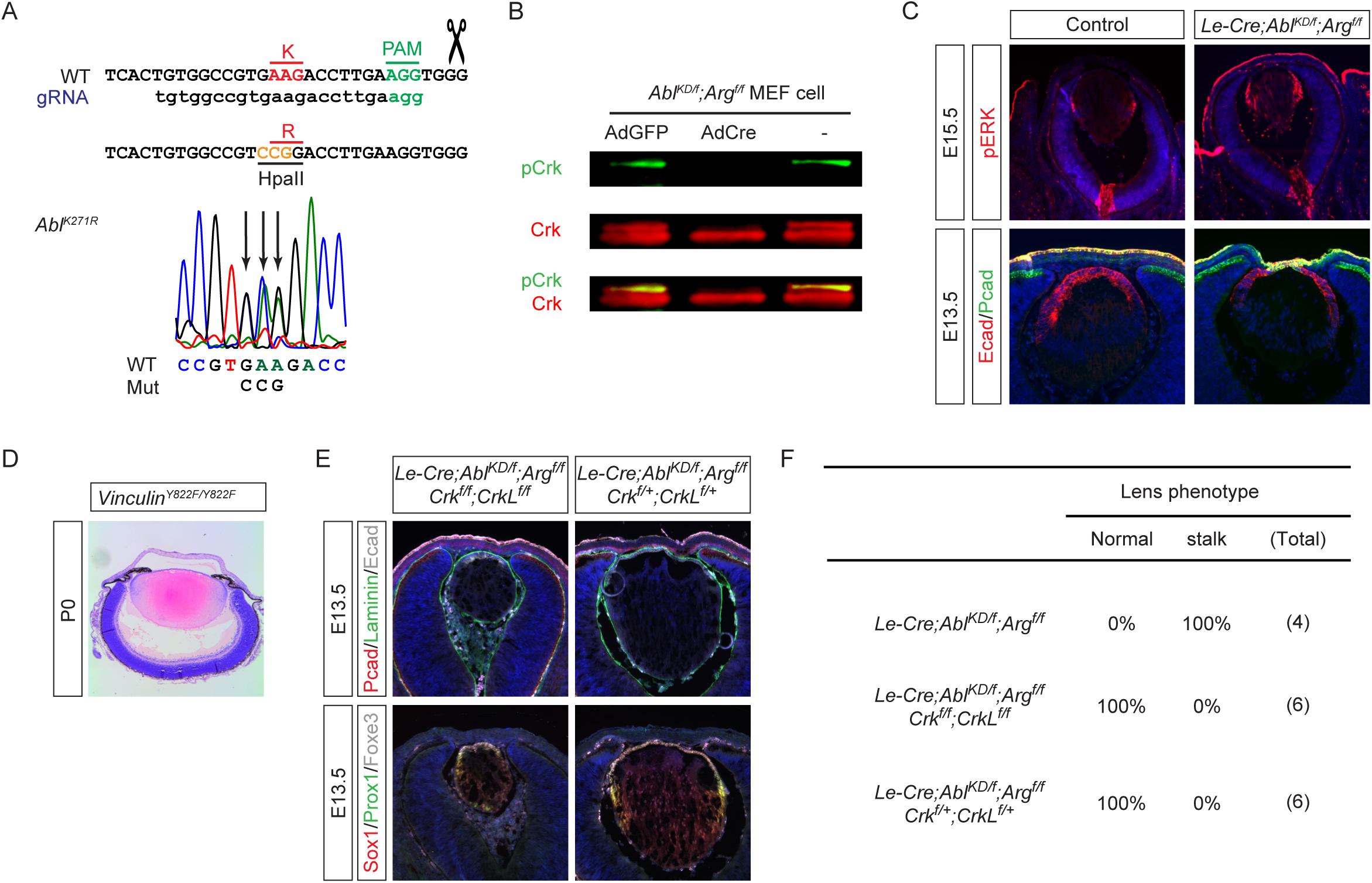
Abl kinase activity is required for lens vesicle closure. **(A)** Schematic representation of the CRISPR-Cas9-mediated generation of the Abl K271R kinase-dead mutation (*Abl^KD^*). **(B)** Western blot analysis demonstrating the loss of pCrk in *Abl^KD^* mutants, confirming the absence of Abl catalytic activity. **(C)** Comparative analysis of lens development showed that *Abl^KD^* phenocopies the *Abl* null lens vesicle closure defect. **(D)** Mutating the Abl phosphorylation site in Vinculin (*Vinculin^Y822F^*) did not affect lens development. **(E)** Genetic interaction study showing rescue of the corneal-lenticular adhesion phenotype in *Abl^KD^* mutants by reducing the dosage of *Crk* and *CrkL*. **(F)** Quantification of the lens stalk phenotype.

Vinculin, which is important for mechanotransduction at adherens junctions, can also be phosphorylated by Abl at Y822(Bays et al., 2014). We generated a Vinculin Y822F mutant (*Vinculin^Y822F^*, details to be published separately) to test whether Abl-mediated regulation of Vinculin activity could be responsible for the lens separation phenotype. Histological analysis of P0 homozygous mutants, however, revealed normal separation of the lens from the cornea, demonstrating that disruption of Abl-mediated Vinculin phosphorylation does not lead to Peters anomaly (Fig. 5D). Lastly, we examined the genetic interaction between *Abl^KD^* mutation and *Crk* deficiency. Similar to the *Abl* conditional mutants, deletion of two to four *Crk* and *CrkL* alleles was sufficient to rescue the lens separation defect in *Le-Cre;Abl ^f/KD^;Arg ^f/f^* mutants (Fig. 5D and E). These findings collectively demonstrate while direct phosphorylation of Crk and Vinculin by Abl is dispensable, the overall kinase activity of Abl plays a crucial role in regulating Crk-mediated signaling pathways during lens morphogenesis.

### Abl kinases stabilize Ptpn12 to modulate p130Cas-Crk signaling

Crk proteins bind to phosphotyrosine residues on p130Cas, bridging interactions with downstream molecules C3G, Sos and Dock1 (Birge *et al*., 2009). Considering that Abl kinases may not directly control the adaptor activity of Crk proteins, we explored whether Abl could influence their upstream interactor p130Cas. To this end, we deleted *Abl/Arg* genetically in MEF cells, which resulted in an increase in p130Cas phosphorylation (Fig. 6A), a finding corroborated in vivo where *Abl/Arg* mutant lenses exhibited enhanced p-p130Cas staining (Fig. 6B, arrowheads). To ascertain if this effect was dependent on Abl ’s kinase activity, we applied pharmacological inhibitors. The dual Src/*Abl* inhibitor Dasatinib reduced phosphorylation of both Src and Crk, diminishing Crk-p130Cas binding as determined by immunoprecipitation. Conversely, the Abl-specific inhibitor Imatinib increased p130Cas phosphorylation without affecting pSrc, leading to enhanced p130Cas-Crk binding (Fig. 6C).

**Figure 6.**
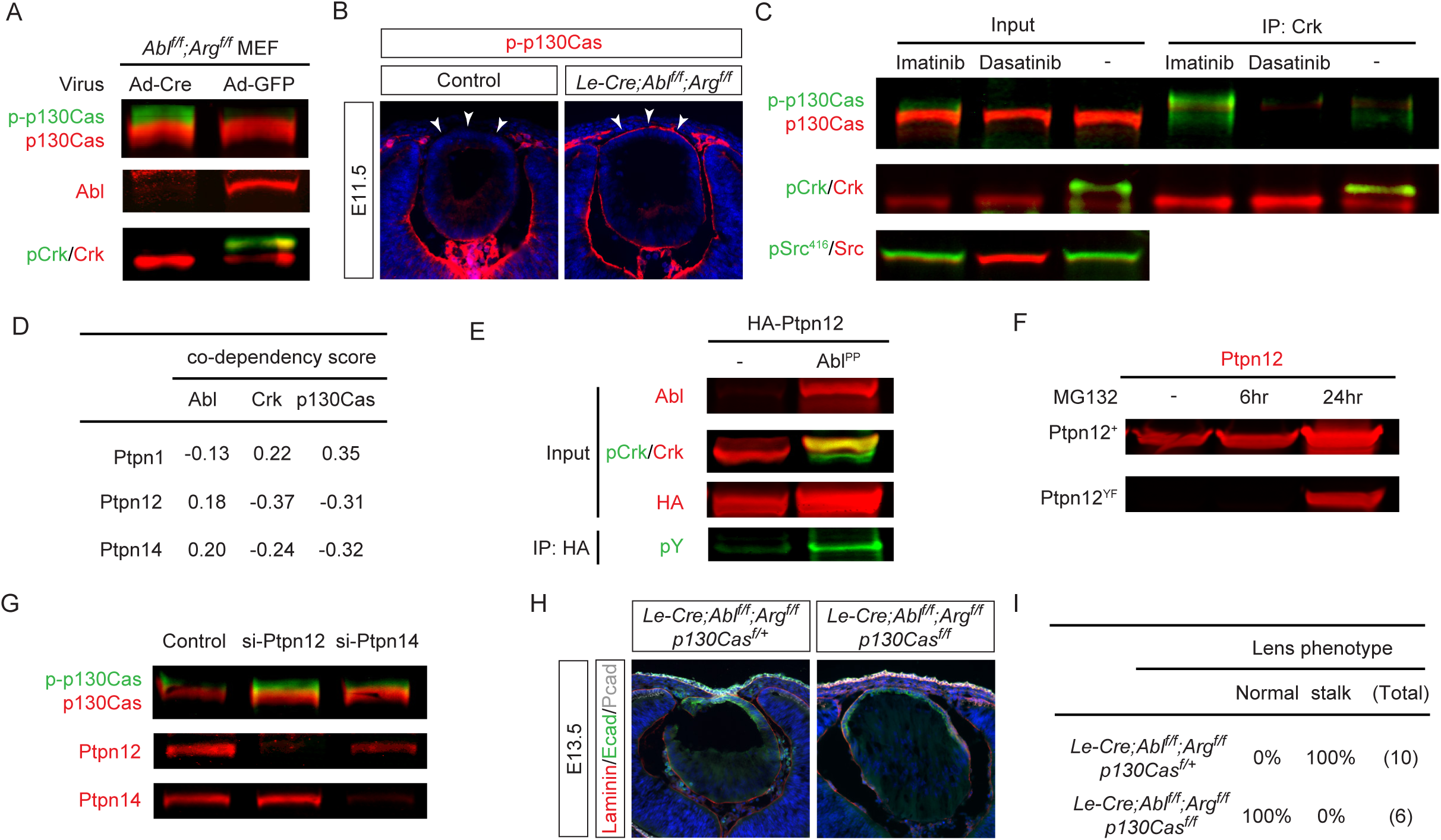
Abl kinases Stabilize Ptpn12 to Regulate p130Cas Phosphorylation. **(A)** Deletion of *Abl/Arg* in MEF cells led to an increased phosphorylation of p130Cas. **(B)** Elevated expression of phosphorylated p130Cas was observed in *Abl/Arg* mutant lenses (arrowheads). **(C)** Inhibition of Abl kinase activity in NIH3T3 cells by Imatinib led to increased phosphorylation of p130Cas and enhanced Crk-p130Cas interaction, as demonstrated by co-immunoprecipitation. **(D)** Analysis from the DepMap database indicates positive co-dependency between Ptpn12 and Abl and negative co-dependency with Crk and p130Cas. **(E)** Expression of constitutively active *Abl^PP^* in HEK293 cells led to tyrosine phosphorylation of Ptpn12 and an increase in its expression level. **(F)** Mutating of tyrosine residues in Ptpn12 significantly reduced its expression, which was mitigated by proteosome inhibitor MG132. **(G)** Increased p-p130Cas levels were observed following siRNA-mediated knockdown of Ptpn12, but not Ptpn14. **(H)** Deletion of p130Cas in *Abl/Arg* mutants rescued the lens vesicle separation phenotype. **(I)** Quantification of the lens stalk phenotype.

Given that *Abl* kinase inhibition increased p130Cas phosphorylation, the most parsimonious interpretation suggests a direct action through a phosphatase. Analysis of the DepMap database revealed three protein tyrosine phosphatases displaying codependency with *Abl*, *Crk*, and *p130Cas* (Fig. 6D). Among them, *Ptpn1* exhibited a negative correlation with *Abl* and a positive correlation with *Crk* and *p130Cas*, making it an unlikely mediator of Abl kinase’s suppression of p130Cas-Crk signaling. Instead, we focused on *Ptpn12* and *Ptpn14*, both positively correlated with *Abl* and previously implicated in p130Cas dephosphorylation (Garton et al., 1996; Zhang et al., 2013). Overexpression of the constitutively active Abl^PP^ not only led to hyperphosphorylation of pCrk as expected but also resulted in tyrosine phosphorylation and increased level of Ptpn12 (Fig. 6E). This is interesting because Ptpn12, also known as PTP-PEST, contains a PEST domain involved in protein degradation which can be regulated by tyrosine phosphorylation (Meyer et al., 2011). Indeed, mutation of all tyrosine residues within Ptpn12 drastically reduced its level, an effect reversible by the proteasome inhibitor MG132 (Fig. 6F), indicating that Abl kinase phosphorylation governs Ptpn12 stability.

Functional assays further confirmed the role of Ptpn12 in controlling p130Cas phosphorylation. siRNA-mediated knockdown of *Ptpn12* and *Ptpn14* efficiently reduced their protein expressions (Fig. 6G) but only *Ptpn12* siRNA increased p130Cas phosphorylation. In vivo, while removing a single copy of *p130Cas* did not affect lens-cornea attachment in *Arg/Abl* mutants, deleting both copies completely blocked persistent attachment (Fig. 6H and I). These results demonstrate that Ptpn12-mediated p130Cas phosphorylation is responsible for Abl kinase regulation of Crk signaling.

### FGF-Abl-Crk signaling regulates the RhoA-Rac1 balance to promote tension-mediated separation of lens vesicle

The lens vesicle and the surface ectoderm are distinguished by the expression of N-and P-cadherin, respectively, while both share E-cadherin expression (Xu *et al*., 2002). Previous cell culture studies have indicated that inhibition of Abl kinases can cause cadherin junction instability (Li and Pendergast, 2011; Zandy et al., 2007). However, examination of *Abl/Arg* mutant lens did not reveal a reduction in cadherin levels in vivo (Fig. 1F and J). Additional deletion of *Cdh1* (encoding E-cadherin) also did not exacerbate the persistent lens stalk phenotype observed in *Abl/Arg* mutants (Fig. 7A). Even simultaneous deletion of both *Cdh2* (encoding N-cadherin) and *Cdh3* (encoding P-cadherin) in the lens vesicle and surface ectoderm did not produce a lens stalk phenotype (Fig. 7B).

**Figure 7.**
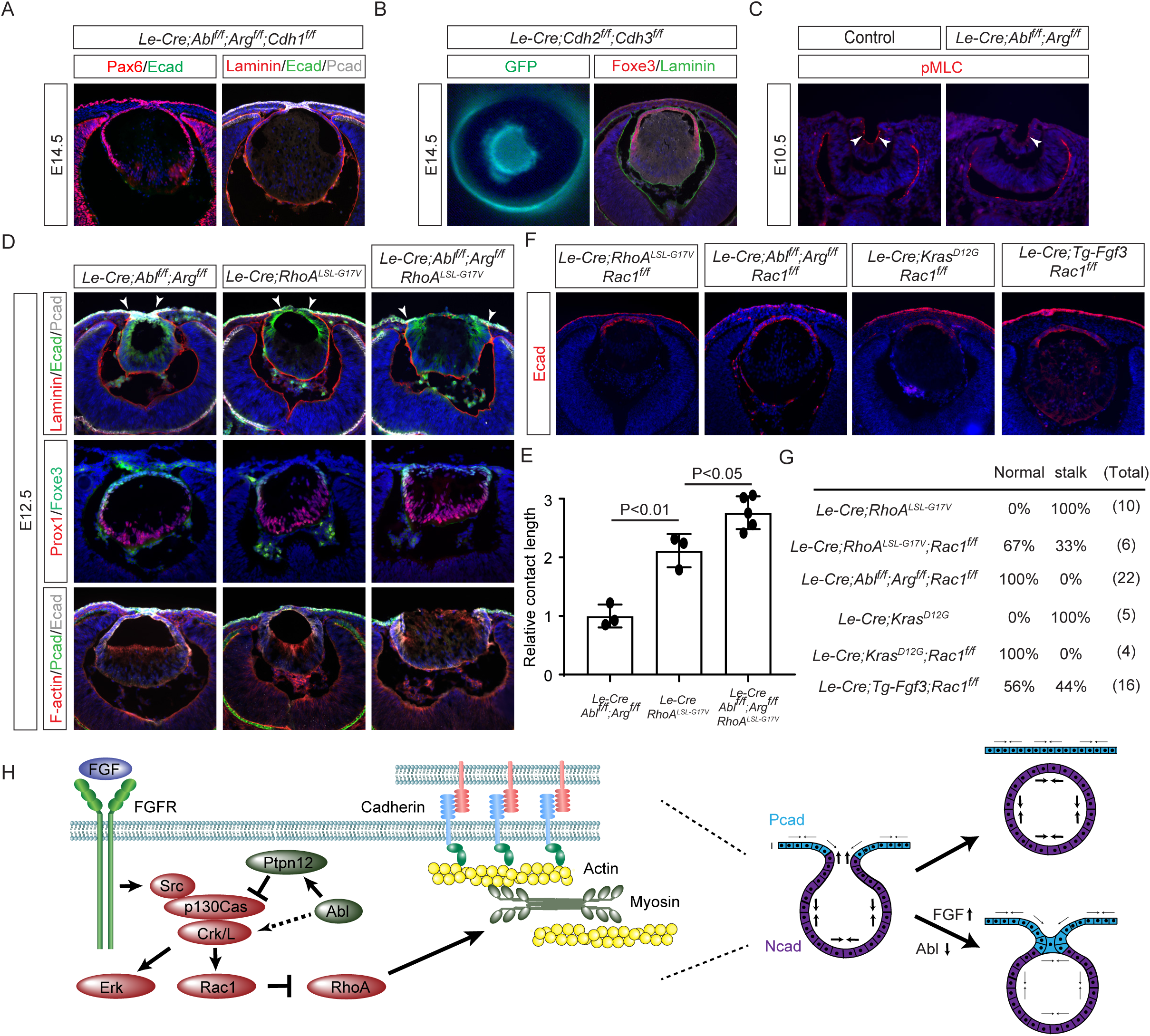
FGF and Abl signaling regulates lens vesicle separation via the p130Cas-Crk-Rac1-RhoA axis. **(A)** *Cdh1* deletion did not exacerbate the lens vesicle separation defect in *Abl/Arg* mutants. **(B)** Combined ablation of *Cdh2* and *3* did not disrupt lens development. **(C)** Phosphorylated MLC (pMLC) was downregulated in *Abl/Arg* mutant lens vesicles (arrowheads). **(D)** The corneal-lens attachment phenotype was similar in *Abl/Arg* and *RhoA^LSL-G17V^*single mutants, but was significantly more severe in the double mutants (arrowheads). **(E)** Quantification of the corneal-lens contact length. One-way ANOVA n=3 for *Abl/Arg* and *RhoA^LSL-G17V^* single mutants, n=5 for double mutants. **(F)** Rac1 deletion rescued lens vesicle separation in *RhoA^LSL-G17V^*, *Abl/Arg*, *Kras^G12D^* and *Tg-Fgf3* mutants. **(G)** Quantification of the lens stalk phenotype. **(H)** The model FGF and Abl kinase signaling in lens vesicle development. FGF and Abl kinase signaling regulates the p130Cas via Src phosphorylation and Ptpn12 dephosphorylation, respectively. This modulation affects the recruitment of Crk and CrkL to p130Cas and the activation of Erk and Rac1, the latter antagonizes RhoA activity. RhoA regulates actomyosin contractility, crucial for separating the lens vesicle from the surface ectoderm.

Although junctional instability caused by Abl kinase inhibition appears to be restricted to in vitro cell culture, it may be an indication of dysregulation of Rho GTPase (Li and Pendergast, 2011; Zandy et al., 2007). Interestingly, we observed that phosphorylation of myosin light chain (pMLC), a key regulator of actinomyosin contractility downstream of RhoA signaling, is significantly weaker in the surface ectoderm than the invaginating lens vesicle (Fig. 7C), suggesting that these two tissues experience differential tension. Importantly, pMLC was significantly reduced in *Abl/Arg* mutant, suggesting that RhoA signaling may be downregulated. To assess the functional significance of RhoA signaling, we employed *Le-Cre* to express a dominant-negative allele of *RhoA* (*RhoA^LSL-G17V^*) (Cortes et al., 2018) and found that it similarly led to the attachment of the lens vesicle to the ectoderm (Fig. 7D). Notably, combining *RhoA^LSL-G17V^* with *Abl/Arg* knockout extended the attachment interface significantly more than either mutation alone (Fig. 7E), indicating a genetic interaction between *RhoA* and *Abl* in facilitating lens vesicle separation. Considering Crk’s role in recruiting the guanine exchange factor (GEF) to activate Rac1 (Birge *et al*., 2009), which is known to have an antagonistic relationship with RhoA (Chauhan et al., 2011), we hypothesized that defective Abl-Crk signaling may lead to RhoA downregulation as a result of increased Rac1 activity. This model predicts that deletion of Rac1 should rescue the *RhoA^LSL-G17V^*lens phenotype. Indeed, we found that *RhoA^LSL-G17V^/Rac1* double mutant reversed the lens stalk phenotype (Fig. 7F-G). Moreover, *Rac1* deletion alone sufficed to prevent the lens stalk in *Abl/Arg* mutants and ameliorated the phenotype in *Le-Cre; Kras ^G12D^* and *Tg-Fgf3* mutants. These results demonstrated that the RhoA-Rac1 balance mediates FGF-Abl-Crk signaling in lens vesicle separation.

## Discussion

In this study, we uncovered a previously unrecognized role for Abl kinases in modulating FGF signaling. Our findings demonstrate the remarkable potency of Abl kinases in FGF signaling as *Abl/Arg* deficiency can partially restore lens development even in the absence of FGF signaling. However, loss of Abl kinases also led to persistent corneal-lenticular adhesion, a hallmark of Peters anomaly in humans. This phenomenon is not caused by elevated ERK signaling. Instead, Abl kinases are crucial for downregulating Crk activity to prevent Peters anomaly (Fig. 7H). Contrary to long-standing beliefs, our data show that while Abl kinases can readily phosphorylate Crk and CrkL, they are gratuitous phosphorylation functionally inconsequential for Crk and CrkL protein activity. Rather, Abl kinases enhance the stability of Ptpn12 through tyrosine phosphorylation, which in turn promotes the dephosphorylation of p130Cas and its interaction with Crk. This cascade of events suppresses Rac1 signaling while enhancing RhoA activity, ultimately stimulating actinomyosin contractility essential for closing the lens vesicle and detaching it from the surface ectoderm. Our findings position Abl kinases as critical regulators in the interplay between FGF signaling and tension-mediated separation of the lens vesicle. This work not only advances our understanding of Abl kinases but also provides insights into the molecular mechanisms underlying Peters anomaly, potentially opening new avenues for therapeutic interventions.

Despite the success of Abl inhibitor Imatinib (Gleevac) for treating chronic myeloid leukemia (CML), the mechanism of Abl kinases in normal physiological conditions is far from understood (Greuber *et al*., 2013). Extensive research has established that tyrosine phosphorylation of Crk proteins is essential for their functionality (Birge *et al*., 2009; Bradley and Koleske, 2009). However, these studies typically utilized overexpressed *Crk* mutants in vitro. Using a genetic knock-in strategy, we introduced phosphorylation-deficient *Crk* and *CrkL* mutants at native expression levels, effectively eliminating interference from wild type *Crk* genes. This approach rigorously tested whether the Abl kinase phosphorylation sites are essential for Crk protein function in living organisms. Despite the sophisticated autoinhibitory structure of phosphorylated Crk (Jankowski *et al*., 2012; Saleh *et al*., 2016), our genetic data demonstrate that these Abl kinase phosphorylation events do not alter function under normal physiological conditions. This underscores the necessity of in vivo studies to determine the real-world relevance of in vitro interactions and prompts a reevaluation of molecular mechanisms previously believed to depend on Crk phosphorylation, including the oncogenic mechanism of v-Crk. Instead, our findings indicate that Abl kinases indirectly reduce Crk signaling by stabilizing the p130Cas phosphatase Ptpn12, which features a c-terminal PEST domain known for signaling protein degradation. Previous studies have also suggested that tyrosine phosphorylation of the PEST domain may influence protein stability (Meyer *et al*., 2011). Ptpn12 has a wide range of physiological functions. Future research should focus on whether Abl kinase phosphorylation serves as a general mechanism for regulating Ptpn12.

According to the classic Differential Adhesion Hypothesis (DAH), cell sorting and tissue segregation are driven by variations in adhesion strength, primarily mediated by cadherins through differential homophilic and heterophilic interactions (Steinberg, 1963). At first glance, this model appears readily applicable to lens development, as the ectoderm expresses P-caderin and the lens expresses N-cadherin, despite both sharing Ecad (Fig. 7K). However, our findings indicate that the genetic ablation of both P-caderin and N-cadherin during lens development does not result in a discernible phenotype, suggesting that differential cadherin expression may not be crucial for the segregation of the lens vesicle from the surface ectoderm. These results stand in contrast to previous studies demonstrating that the combined loss of N-caderin and E-caderin leads to a persistent lens stalk (Pontoriero *et al*., 2009). These observations align with a recent revision of the DAH, which emphasizes the critical role of cortical tension driven by actomyosin contraction rather than mere cadherin-mediated adhesion (Winklbauer, 2015). In this updated model, cadherins primarily function to mechanically link neighboring cells, thereby allowing differential cortical tension to dictate cell sorting and tissue segregation. During lens development, the lens vesicle cells undergo apical constriction and experience significantly higher tension than the surrounding surface ectoderm, as indicated by the elevated expression of phosphorylated myosin light chain (pMLC). We propose that this disparity in tissue tension not only facilitates the invagination of the lens vesicle but is also crucial for its ultimate separation from the surface ectoderm (Fig. 7K). In contrast, *Abl/Arg* deficiency leads to reduced actomyosin contractility, primarily through decreased RhoA signaling, resulting in insufficient tissue tension to effectively segregate the lens vesicle from the surface ectoderm. These results highlight the crucial role of Abl kinases in regulating cortical tension for tissue self-organization.

Peters anomaly, the predominant cause of congenital corneal opacity, presents significant challenges in pediatric ophthalmology (Bhandari *et al*., 2011). While corneal transplantation is feasible in young children, it is associated with a high failure rate, underscoring the need for alternative therapeutic approaches. This condition has a diverse genetic etiology, primarily involving transcription factors that are challenging to target. Although *FGFR2* has been implicated in Peters anomaly, the molecular mechanisms remain unclear (McCann *et al*., 2005; Okajima *et al*., 1999). Our study links FGF signaling to the Abl-Ptpn12-p130Cas-Crk pathway, which in turn modulates Rho GTPase activity during lens vesicle separation. This identifies multiple potential targets within this pathway, including kinases, phosphatases, and Rho GTPase. Our research underscores the critical role of tissue tension in ocular development and suggests that its modulation could be a key mechanism in the pathogenesis of Peters anomaly. This insight opens the possibility that targeting this tension-mediated mechanism could emerge as a therapeutic strategy for managing Peters anomaly and potentially other related ocular developmental disorders.

## Methods and Materials

### Mice

*Abl^flox^* and *Arg^flox^* mice have been previously reported (Kim et al., 2019; Moresco et al., 2005). *Le-Cre* mice are obtained from Richard Lang (Children’s Hospital Research Foundation, Cincinnati, OH) (Ashery-Padan *et al*., 2000), *Cdh2^flox^* from Yulia Komarova (University of Illinois College of Medicine, Chicago, IL) (Kostetskii et al., 2005), *Crk^flox^* and *CrkL^flox^* from Dr. Tom Curran (The Children’s Research Institute, Children’s Mercy Kansas City), *Erk1^KO^*, *Erk2^flox^* from Jian Zhong (Weill Cornell Medicine, White Plains, NY), *Fgf3^OVE391^* from Dr. Michael Robinson, Miami University, Oxford, OH) (Robinson et al., 1998), *Fgfr2^flox^* from Dr. David Ornitz (Washington University Medical School, St Louis, MO) (Yu et al., 2003). *p130Cas^flox^* from Dr. Martin Riccomagno (University of California, Riverside, CA) (Riccomagno et al., 2014), *Rac1^flox^* from Dr. Feng-Chun Yang (Indiana University School of Medicine) (Glogauer et al., 2003), *RhoA^LSL-G17V^* mice from Teresa Palomero (Columbia University, New York, NY) (Cortes *et al*., 2018), *LSL-Kras^G12D^* from the Mouse Models of Human Cancers Consortium (MMHCC) Repository at National Cancer Institute (Tuveson et al., 2004). *Cdh1^flox^* (Stock No: 005319) and *Cdh3^KO^* (Stock No: 003180), *Fgfr1^flox^*(Stock No: 007671), *Mek1 ^DD^*allele (Stock No: 012352) mice were obtained from Jackson Laboratory. *Abl^K271R^*, *Crk^Y221F^* (*Crk^YF^*), and *CrkL^Y207F^* (*CrkL^YF^*) were generated at the Columbia University Transgenic Core Facility. Animals were maintained in a mixed genetic background, and at least three animals were analyzed for each genotype. In all conditional knockout experiments, mice were maintained on a mixed genetic background and *Le-Cre* only or *Le-Cre* and heterozygous flox mice were used as controls. All animal experiments were performed according to protocols approved by the Columbia University Institutional Animal Care and Use Committee.

### Histology, Immunohistochemistry, and Immunocytochemistry

Hematoxylin and Eosin (H&E) staining was conducted on the paraffin-embedded tissue sections. Immunohistochemistry (IHC) was performed on both paraffin and cryo-sections, following previously established protocols (Carbe et al., 2012; Carbe and Zhang, 2011). For the detection of pErk (Cell Signaling, #4370) and pP130Cas (Cell Signaling, #4011), signal amplification was achieved using the Tyramide Signal Amplification Kit (PerkinElmer Life Sciences, Waltham, MA). Additional antibodies used included: Cleaved caspase3 (#9662), E-cadherin (#3195), N-cadherin (#13116) (all from Cell Signaling), Ki67 (BD Pharmingen, #550609), Foxe3 (sc-377465), Jag1(#6011), Maf (#7866) (all from Santa Cruz), Laminin (Sigma, #L9393), p57 (#75947) (from Abcam), Pax6 (Biolegend, #PRB-278P), pHH3 (Millipore, #06-570), and Prox1 (Covance, PRB-238C). Antibodies against α-and γ-crystallins were kindly provided by Sam Zigler (National Eye Institute, Bethesda, MD).Alexa Fluor 488 and 568 Phalloidin were sourced from ThermoFisher (#A12379 and #A12380). TUNEL staining was performed using the TUNEL Enzyme (Sigma, #11767305001) and TUNEL Label Mix (Sigma, #11767291910).

### Cell Culture

Primary mouse embryonic fibroblasts (MEFs) were isolated from E13.5 embryos and cultured in Dulbecco’s Modified Eagle’s Medium (DMEM) supplemented with 10% fetal bovine serum (FBS), following established protocols (Qu et al., 2011). NIH3T3 and HEK293 cells were also maintained in DMEM with 10% FBS. Cells were starved for 24 hours before treatment with Imatinib (200 μM; Santa Cruz, #sc-202180A) or Dasatinib (1 nM; Santa Cruz, #sc-358114) for one hour prior to harvesting. HEK293 cells were treated with MG132 (20 μM, Sigma, M7449) up to 24 hours.

### Transfection and Transduction

For gene deletion, MEF cells carrying homozygous floxed alleles were infected overnight with Ad5CMV-eGFP or Ad5CMVCre-eGFP (Gene Transfer Vector Core, University of Iowa, IA) at a multiplicity of infection of 500 plaque-forming units per cell in DMEM. The cells were then cultured for an additional five days. For plasmid transfection, NIH3T3 and HEK-293 cells were transfected with Lipofectamine 3000 (ThermoFisher, #L3000015) following the manufacturer’s guidelines. Plasmid amounts were based on a scaling table. The plasmids were pcDNA3.1 c-Abl^pp^ (Dr. Michael K. Lee, University of Minnesota) (Karim et al., 2020), pCMV3-N-HA-PTPN12 (Sino Biological, #HG11556-NY), and pTwist-CMV-PTPN12YF synthesized by Twist Bioscience, with all tyrosine residues in PTPN12 replaced by phenylalanine and inserted into the pTwist-CMV plasmid. For siRNA transfection, NIH3T3 and HEK-293 cells were plated at 50% confluency. Cells were transfected with 100 nM of siRNAs targeting PTPN12, PTPN12 Smart Pool, PTPN14, and GAPDH (Dharmacon, #J-042199-05-0002, #L-042199-00-0005, #J-040086-05-0002, and #D-001830-20-05, respectively), using Lipofectamine 3000. To enhance transfection efficiency, the transfection procedure was repeated 24 hours later.

### Immunoprecipitation and Western Blot

After the designated treatments, cells were washed with warm phosphate-buffered saline (PBS) and stimulated with pervanadate for 2 minutes. Following stimulation, cells were rinsed in cold PBS and lysed using ice-cold CelLytic Reagent (Sigma, #C2978) supplemented with a protease inhibitor cocktail (Thermo Fisher, #78841). Antibodies were immobilized on Dynabeads (ThermoFisher, #10002 and #10004) and incubated with the cell lysate for immunoprecipitation. The resulting precipitates were dissolved in SDS sample buffer. Samples were denatured at 95°C for 5 minutes before being loaded onto SDS–polyacrylamide gel electrophoresis (SDS-PAGE) gels for separation. Phos-tag acrylamide (FUJIFILM Wako Chemicals, NC0232095) was incorporated into gels to achieve high resolution separation of phosphorylated and phospho-deficient Crk/CrkL. The following antibodies were used for immunoblotting: pErk1/2 (Santa Cruz, #sc-7383), Erk1/2 (Cell Signaling, #4695), Abl (BD Pharmingen, #554148), Crk (BD Pharmingen, #610036), pCrkII-Tyr221 (Cell Signaling, #3491), pCrkL-Tyr207 (Cell Signaling, #34940), GFP (Aves Labs, #GFP-1010), HA (Cell Signaling, #2367 and #3274), P130Cas (BD Pharmingen, #610271), pP130Cas-Tyr165 (Cell Signaling, #4011), PTP-PEST (Santa Cruz, #sc-65229), PTPN14 (Cell Signaling, #13808), and phospho-tyrosine (Cell Signaling, #9411).

### Quantification and Statistical Analysis

Cell proliferation and apoptosis were assessed by calculating the ratios of Ki67-, pHH3-, and TUNEL-positive cells to DAPI-positive cells. The length of the lens vesicle’s contact with the surface ectoderm, along with lens size, were quantified using ImageJ software from the NIH. The number of lens stalks was determined through both whole mount eye imaging and immunohistochemistry. Statistical analyses were conducted using t-tests for pairwise comparisons and one-way ANOVA for comparisons involving three or more groups

## Acknowledgements

The authors thank Drs. Ruth Ashery-Padan, Tom Curran, Yulia Komarova, Richard Lang, Michael K. Lee, David Ornitz, Martin Riccomagno, Michael Robinson and Jian Zhong for mice and reagents. The work was supported by grants from NIH (R01EY017061 and R01EY025933 to X.Z.). Q.W. is supported by a Pathway to Independence Award (K99EY032171). The Columbia Ophthalmology Core Facility is supported by NIH Core grant 5P30EY019007 and unrestricted funds from Research to Prevent Blindness (RPB).

